# Two-Metal Ion Mechanism of DNA Cleavage by Activated, Filamentous SgrAI

**DOI:** 10.1101/2024.05.01.592068

**Authors:** Zelin Shan, Andres Rivero-Gamez, Dmitry Lyumkis, N. C. Horton

## Abstract

Enzymes that form filamentous assemblies with modulated enzymatic activities have gained increasing attention in recent years. SgrAI is a sequence specific type II restriction endonuclease that forms polymeric filaments. SgrAI filamentation increases enzymatic activity by up to three orders of magnitude and additionally expands its DNA sequence specificity. Prior studies have suggested a mechanistic model linking the structural changes accompanying SgrAI filamentation to its accelerated DNA cleavage activity. In this model, the conformational changes that are specific to filamentous SgrAI maximize contacts between different copies of the enzyme within the filament and create a second divalent cation binding site in each subunit, which in turn facilitates the DNA cleavage reaction. However, our understanding of the atomic mechanism of catalysis is incomplete. Herein, we present two new structures of filamentous SgrAI solved using cryo-electron microscopy (cryo-EM). The first structure, resolved to 3.3 Å, is of filamentous SgrAI containing an active site mutation that is designed to stall the DNA cleavage reaction, which reveals the enzymatic configuration prior to DNA cleavage. The second structure, resolved to 3.1 Å, is of WT filamentous SgrAI containing cleaved substrate DNA, which reveals the enzymatic configuration at the end of the enzymatic cleavage reaction. Both structures contain the phosphate moiety at the cleavage site and the biologically relevant divalent cation cofactor Mg^2+^ and define how the Mg^2+^ cation reconfigures during enzymatic catalysis. The data support a model for the activation mechanism that involves binding of a second Mg^2+^ in the SgrAI active site as a direct result of filamentation induced conformational changes.

## Introduction

Enzyme filamentation involves the polymerization of multiple copies of the same protein into long linear, helical, or tubular structures. Filamentation has been observed for decades, yet only recently has it been generally acknowledged to be a widespread mechanism of enzyme regulation (1–5). To date, over 30 enzymes from diverse biochemical pathways and from organisms representing all domains of life have been shown to form filamentous assemblies (1,2). A subset of these enzymes have been subjected to detailed studies to determine how filamentation affects enzyme activity (6,7). Many of the same enzymes known to form filaments *in vitro* also form large self-assemblies in cells that are visible by fluorescence microscopy, also known as cytoophidia (1,2,5,8–13). In some cases, a direct relationship between cytoophidia formation and enzyme polymerization/filamentation has been shown (5,14). The purpose of these cellular substructures is not fully understood. There have been several suggested roles. For example, filamentation may lead to the rapid activation/inactivation of cellular enzymes, or alternatively enable control of the levels of active enzyme in the cell. In some cases, the self-assemblies have been suggested to nucleate phase separated bodies containing enzymes from a biochemical pathway to provide greater catalytic throughput within select pathways (12,14–16). Additional possible roles include modulating enzymatic specificity or performing other specialized functions (1,2,7).

One of the best studied enzymes that forms filamentous assemblies is the type II restriction endonuclease SgrAI. SgrAI cleaves an 8 base pair recognition sequence, CR|CCGGYG (R=A or G, Y=C or T, | denotes cleavage site) producing “sticky” ends consisting of a 4 base pair overhang 5’CCGG (6,17). The recognition sequence possesses degeneracy at the second and seventh nucleotide, which lead to a total of 3 different sequences in double-stranded DNA that are known as primary recognition sequences or primary sites. Under conditions that favor enzyme filamentation, SgrAI will also rapidly cleave fourteen additional sequences known as secondary recognition sequences or secondary sites, with the patterns CC|CCGGYG and CR|CCGGYD (D=A, C, or T)(18–20). Binding to primary sites with sufficient base pairs flanking the recognition sequence induces filamentation, and when in the filamentous state, SgrAI cleaves its primary and secondary sequences 200- and 1000-fold faster, respectively, than when in the non-filamentous state (19–21).

Dimeric SgrAI binds both primary and secondary dsDNA recognition sequences in a 1:1 complex known as the DNA bound SgrAI dimer (or DBD) (19,22). The SgrAI filament is a left-handed helix with approximately 4 DBD per turn (**Fig. 1A**) (23). Comparison of DBD conformations in the filamentous and non-filamentous states shows a difference in the positioning of one subunit of the dimer relative to the other that can be characterized as an ∼11° rotation about an axis roughly parallel to the helical axis of the bound DNA **(Fig. 1B**) (24). Inspection of the residues at the dimeric interface shows how conformational changes in the protein accommodate this rotation, which propagate to the active site and shift a segment of SgrAI (residues 184-187) closer to the bound DNA (**Fig. 1C**). The conformational changes were first observed in a cryo-EM structure of SgrAI in the filamentous form bound to a primary site DNA, which was missing the phosphate at the cleavage site (24). However, in this structure, only a single Mg^2+^ ion was observed in the active site in a location known as site A; likewise, only one Mg^2+^ ion was observed in non-filamentous structures of SgrAI DBD bound to DNA (25,26). The shift in residues 184-187 observed within the filamentous structures suggests a mechanism for activation of the DNA cleavage activity wherein a second Mg^2+^ binding site is created. Two Mg^2+^ ions, in sites A & B, respectively (**Fig. 1D**), are predicted through the “two-metal-ion mechanism”, which is believed to be used by many divalent cation-dependent nucleases as well as other phosphoryl transfer enzymes (27–31). In this mechanism, the two divalent metal cations coordinate oxygen ligands derived from both the protein, the DNA, and water molecules, and perform important functions such as activating the nucleophile, positioning reactive groups, stabilizing the leaving group, and stabilizing the transition state (29,30,32,33). In non-filamentous SgrAI structures, typically only site A is occupied. The absence of site B occupancy by Mg^2+^ is thought to be the origin behind the low intrinsic DNA cleavage activity of SgrAI in the non-filamented state (25,26). Notably, the presence of two Mg^2+^ ions in filamentous SgrAI has never previously been observed.

**Figure 1.**
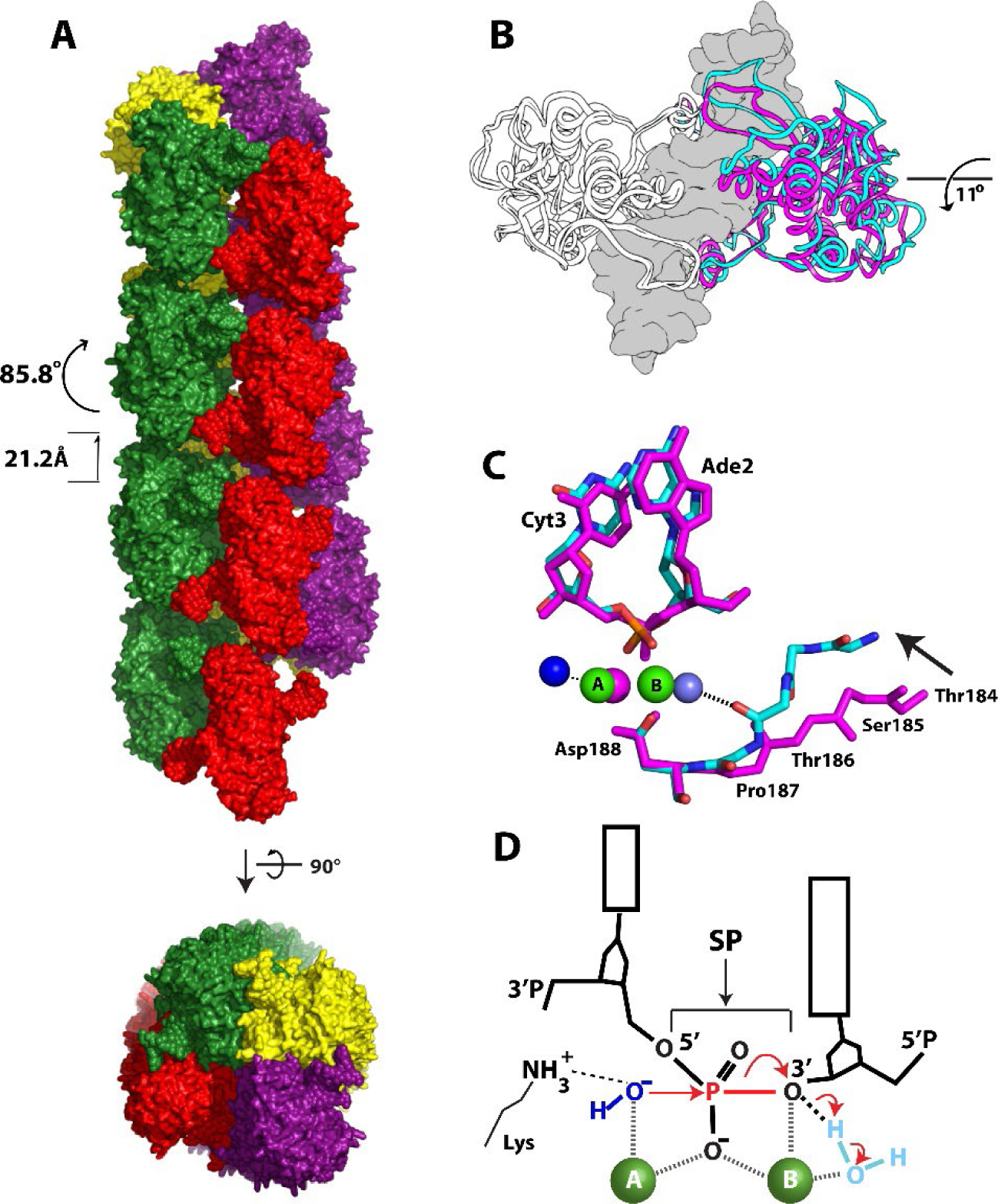
Structures of filamentous and non-filamentous SgrAI and the two metal ion mechanism. **A.** Structure of the SgrAI filament showing approximately four DNA bound SgrAI dimers (DBD, each colored individually in red, green, yellow or purple) per turn in a left-handed helix. **B.** Superposition of filamentous (cyan) and non-filamentous (magenta) SgrAI structures using one chain of a DBD showing an 11° rotation of the other subunit. DNA from the nonfilamentous structure shown in grey in surface rendering. **C.** View of the active site in the superimposed chains from filamentous (cyan, from PDB ID 7SS5) and non-filamentous (magenta, from PDB ID 3DVO) DBD showing a shift of residues 184-187, which creates a second divalent cation binding site in the filamentous structure. **D.** Schematic of the two-metal-ion mechanism. The nucleophile of the reaction is a water or hydroxide (dark blue) which is positioned by its coordination to metal ion A for in-line attack on the phosphorus atom of the scissile phosphate (SP). Both metal ions A and B coordinate a non-bridging oxygen of the SP, and metal ion B coordinates the leaving group as well as a water molecule positioned to donate a proton to the leaving group following bond cleavage (the bond to be cleaved is shown in red). An active site lysine found in many restriction endonucleases is shown interacting with the nucleophile and may serve in its positioning and activation.

Occupation of a site B metal in SgrAI bound to uncleaved DNA was first observed in the structure of filamentous SgrAI bound to a primary site DNA and Ca^2+^ ions (34). However, Ca^2+^ is not the biologically relevant cofactor for SgrAI-mediated cleavage. The divalent cation Ca^2+^ is often used to stall Mg^2+^-dependent DNA cleavage reactions in an attempt to capture the active site structure immediately prior to the DNA cleavage reaction, because Ca^2+^ often binds roughly where Mg^2+^ is expected to bind (35–38). Since Ca^2+^ also inhibits enzymatic cleavage, the Ca^2+^-bound enzymatic state must differ from the Mg^2+^-bound enzymatic state; the underlying basis for the differences between the two ions has been the subject of debate (32,33,38). Ca^2+^ has a larger ionic radius than Mg^2+^ and is less stringent with respect to the coordination geometry of its ligands (39,40). Ca^2+^ ions also induce deprotonation of coordinated water molecules to a lesser degree than Mg^2+^ (the pKa of Ca^2+^ coordinated water is 12.6 while that of Mg^2+^ is 11.4) (41,42). These factors have been suggested to be the source of inhibition of DNA cleavage in some enzymes by Ca^2+^. Thus, although the use of Ca^2+^ can provide insights relevant to enzymes containing Mg^2+^ binding sites, to gain a complete understanding of the enzyme mechanism, it is critical to obtain high-resolution structures with the Mg^2+^ metal ion.

Here, we present two new cryo-EM structures of filamentous SgrAI containing the biologically relevant cofactor Mg^2+^. The first structure, resolved to 3.3 Å, contains the active site mutation K242A, which inhibits DNA cleavage and leaves an intact phosphodiester bond at the cleavage site (hereafter known as the scissile phosphate, or SP). The second structure, resolved to 3.1 Å, is derived from WT SgrAI bound to an intact primary site DNA. This structure contains cleaved DNA in the active site following termination of the cleavage reaction. Both structures show the presence of Mg^2+^ in both sites A and B, thus supporting a model for activation of the DNA cleavage reaction in filamentous SgrAI, which involves the creation of a binding site for a second Mg^2+^ at site B. Based on these data, we propose an updated mechanistic model for filamentation-induced catalytic activity of SgrAI.

## Results

### Overview of SgrAI_K242A_/40-1/Mg^2+^ and SgrAI_WT_/40-1/Mg^2+^ cryo-EM structures

To capture a high-resolution snapshot of enzymatic catalysis, it is necessary to stall the enzyme in a defined state. To determine the enzymatic configuration of SgrAI and the position of preferred metal ion cofactors within the active site at distinct stages of catalysis, we set out to assemble two types of SgrAI filaments, both containing the native Mg^2+^ metal ions. The first assembly employed the mutation K242A, which is predicted to greatly reduce enzymatic activity, but would inform on the position of the Mg^2+^ ions in the step immediately preceding catalysis. The second assembly employed the WT enzyme, which was allowed to catalytically cleave DNA to completion and would thus represent the product form. Both assemblies are expected to be stable, yielding homogeneous complexes for structural studies.

To make the SgrAI_K242A_/40-1/Mg^2+^ filamentous assembly, we incubated K242A SgrAI with a 40 bp DNA containing the primary sequence CACCGGTG (termed 40-1 DNA) in buffer containing MgCl_2_ and incubated for 30 minutes at room temperature prior to cryo-EM grid preparation. We observed filaments on cryo-EM grids and collected a dataset containing 3,488 cryo-EM movies of the sample. Image processing using previously described workflows (24,34) revealed a map resolved to 3.3 Å, which was sufficient for visualization of Mg^2+^ ions. We used a previous model as a starting point (PDB ID 7SS5) (34) to build and refine an atomic model of the filamentous form of K242A SgrAI protein bound to 40-1 DNA, yielding a structure with good geometry and a cross-correlation between map and model of 0.77 (**Table S1**). To make the SgrAI_WT/_40-1/Mg^2+^ filamentous assembly, we mixed WT SgrAI with the same 40-1 DNA and identical buffer conditions but allowed the reaction to proceed for 1.5 hours. This ensured that the cleavage reaction would go to completion. Cryo-EM images again revealed the presence of filaments, and we collected a dataset containing 2,704 movies of the sample. Computational image analysis yielded a map resolved to 3.1 Å (**Table S1**). The refined model again had good geometry and statistics, with a cross-correlation between the map and model of 0.75 (**Table S1**). **Figures S1-S2** show the cryo-EM data, the segmented cryo-EM maps, and refined atomic models for each structure. Finally, **Table S2** shows RMSD of the two new structures to each other and to previously determined structures of SgrAI bound to DNA.

The structure of SgrAI_K242A_/40-1/Mg^2+^ shows that the phosphodiester bond in the mutant form of the protein is intact (**Fig. 2A**). We were able to model two Mg^2+^ ions in the active site, in sites A and B (**Fig. 2A**), and the absence of the K242 side chain is evident (**Fig. 2B**). In contrast, the structure of SgrAI_WT_/40-1/Mg^2+^ shows, as expected, that the SP is now cleaved in the active site (**Fig. 2C**). Two Mg^2+^ions are also evident in the active site, at the positions expected for sites A and B of the two-metal-ion mechanism (**Fig. 2D-E**). A comparison of the active site geometries shows that the position of the Mg^2+^ ions in sites A and B are conserved, although there are slight differences in their coordination, which can be attributed to the nature of the SP (cleaved or uncleaved) and the presence or absence of the K242 side chain, and will be described in more detail below (**Fig. 3A-B**). Collectively, these two structures provide before and after snapshots of the cleavage mechanism in the presence of the biologically relevant cofactor Mg^2+^.

**Figure 2.**
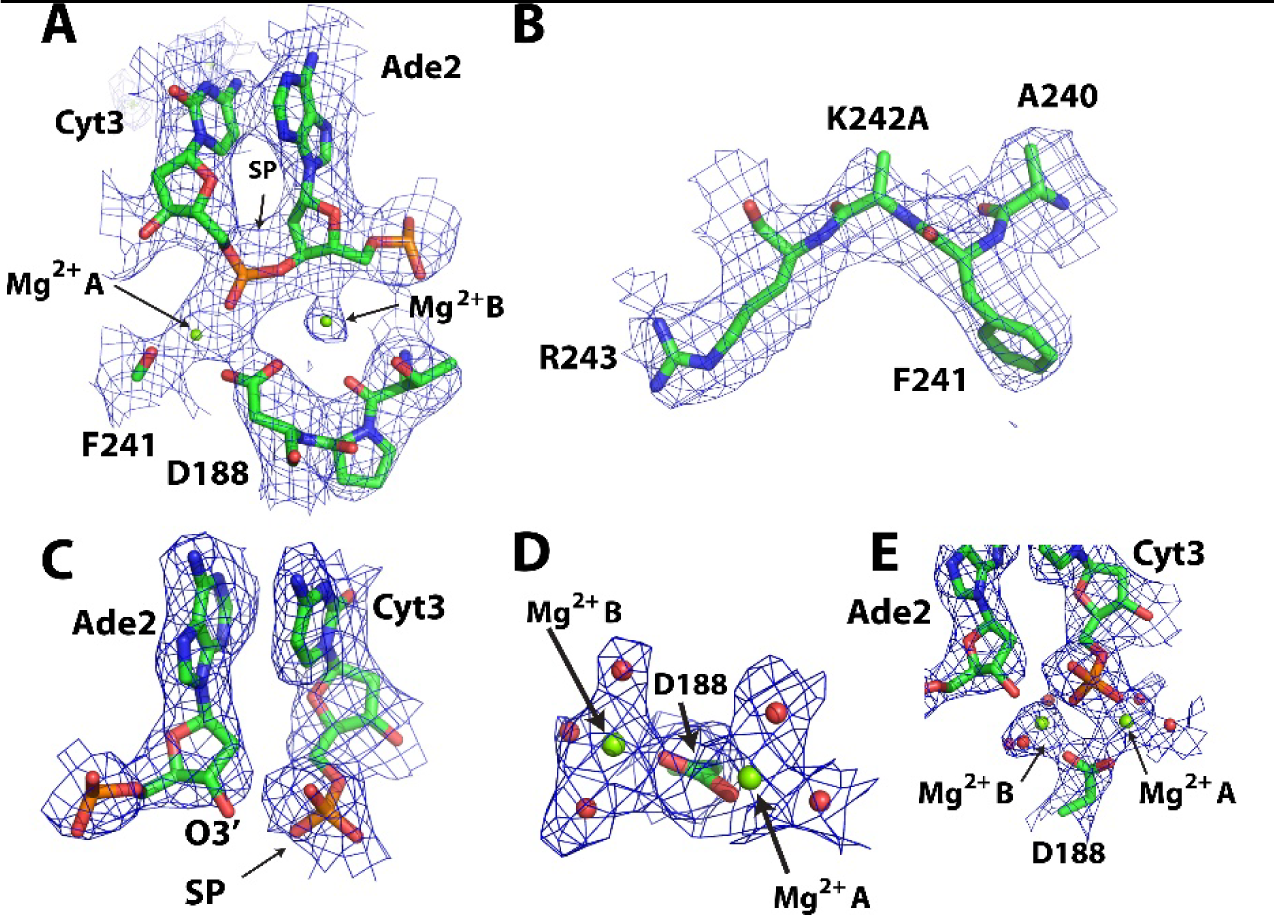
Atomic models and experimental cryo-EM density maps in the active site. SgrAI_K242A_/40-1/Mg^2+^ around active site. Map contoured at 3σ. **B.** As in A, showing area around the K242A mutation. Map contoured at 4σ. **C.** SgrAI_WT_/40-1/Mg^2+^ at scissile phosphate. Map contoured at 4.5σ. **D-E.** As in C, but around Mg^2+^ in the active site. Map contoured at 5σ.

**Figure 3.**
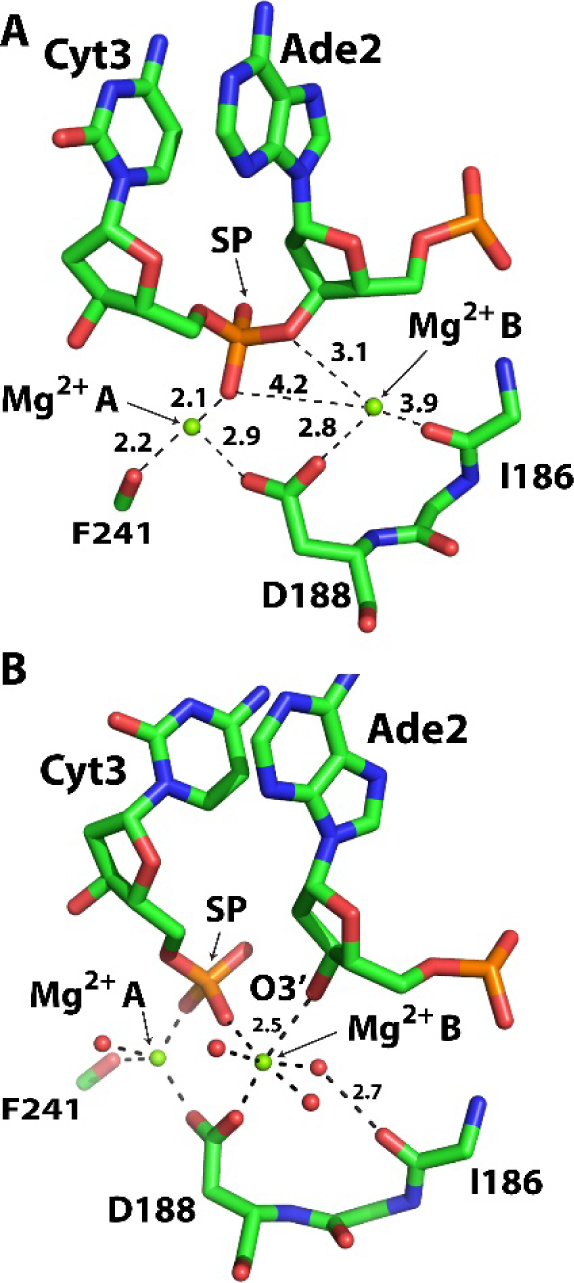
Active site geometry and Mg^2+^ positioning in SgrAI_K242A_/40-1/Mg^2+^ and SgrAI_WT_/40-1/Mg^2+^. **A.** The active site of SgrAI_K242A_/40-1/Mg^2+^. All distances between Mg^2+^ and oxygen ligands are 1.9-2.2 Å unless otherwise indicated. **B.** As in A, but for SgrAI_WT_/40-1/Mg^2+^. Distances shown in Å. SP, scissile phosphate.

### Analysis of SgrAI_K242A_/40-1/Mg^2+^ Structure

To gain insight into the atomic reconfigurations of the enzymatic active site just prior to DNA cleavage, we compared the filamentous SgrAI_K242A_/40-1/Mg^2+^structure to previously determined structures of filamentous SgrAI assemblies. **Figure 4** shows overlays of the active sites of SgrAI_K242A_/40-1/Mg^2+^ with the other previously determined filamentous SgrAI structures, 6OBJ (**Fig. 4A**) and 7SS5 (**Fig. 4B**). The nucleotides align well in these comparisons, but the divalent cations in both sites A and B are significantly displaced, by 1.4 and 1.9 Å, respectively (**Table S3**). The phosphodiester bond of the nucleotide 3′ of the SP (3’P in **Fig. 4A,C**) in SgrAI_K242A_/40-1/Mg^2+^ is similar to all other structures, with the exception of 6OBJ where it rotates to directly coordinate metal ion A. In addition, the conformation of the SP is similar in all structures, with the exception of SgrAI_WT_/40-1/Ca^2+^ (PDB ID 7SS5, slate, **Fig. 4B-C**). The distinct conformation of the SP in 7SS5 appears related to the Ca^2+^ bound in the active site, particularly in site B. The rotation of the SP in 7SS5 allows for both non-bridging oxygens of the SP to each coordinate with the site A and B metal ions. This may result in a configuration that is not conducive for catalysis (**Fig. 4B**), consistent with the absence of DNA cleavage with Ca^2+^ in place of Mg^2+^.

**Figure 4.**
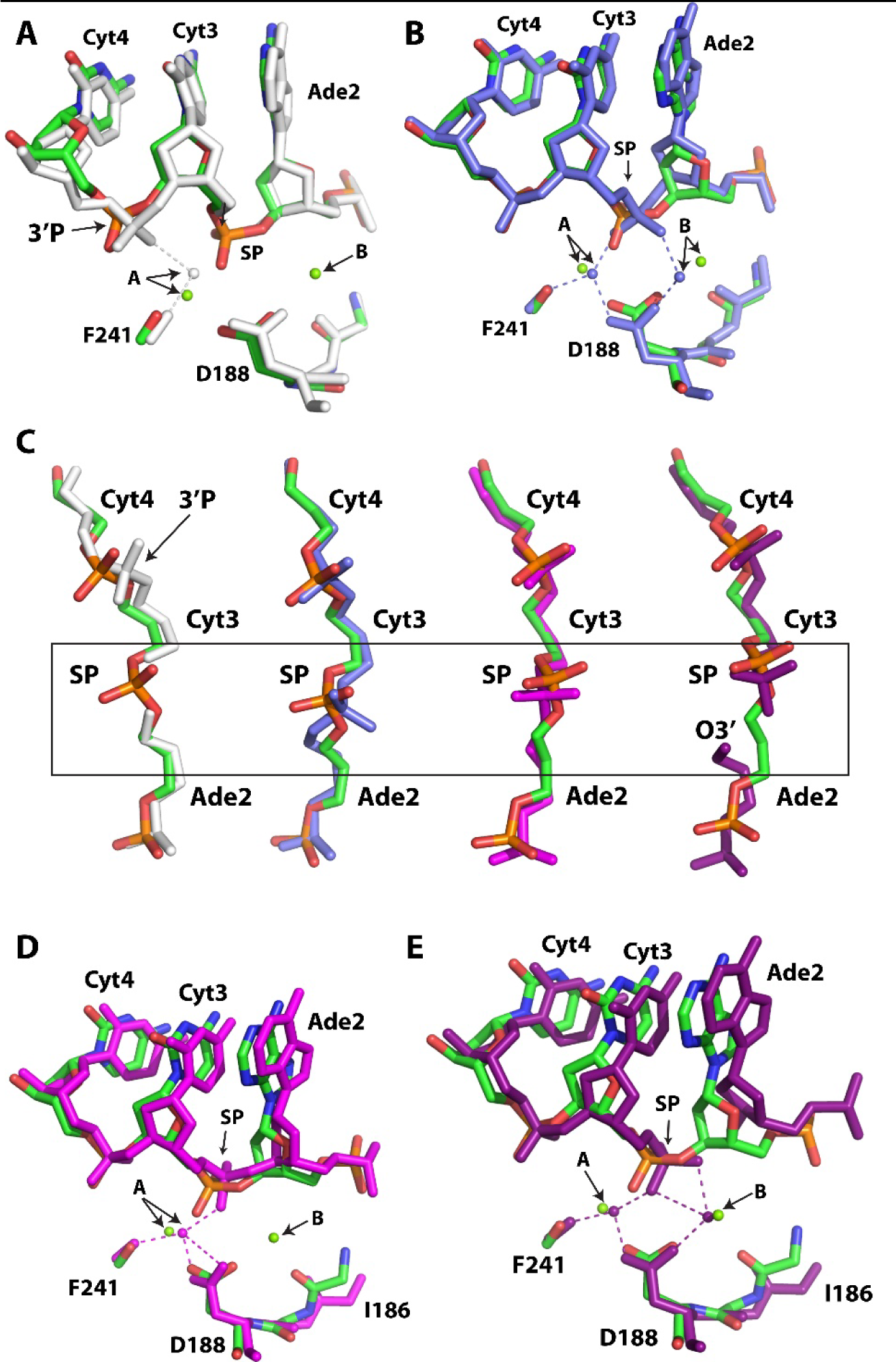
Active site superpositions of SgrAI_K242A_/40-1/Mg with other SgrAI/DNA structures. **A.** Superposition using all atoms of chain A of SgrAI_K242A_/40-1/Mg (green, blue, red, orange) with those of 6OBJ (white). SP, scissile phosphate. **B.** As in A, but with 7SS5 (slate). **C.** As in A, but with view for SP conformation. 3DVO in magenta, 3MQY in dark purple. Boxed region indicates cleavage site. **D.** As in A but with 3DVO (magenta). **E.** As in A but 3MQY (dark purple).

Comparison of SgrAI_K242A_/40-1/Mg^2+^ to non-filamentous structures of wild type SgrAI bound to primary site uncleaved DNA with Ca^2+^ (3DVO, magenta, **Fig. 4D**) or Mg^2+^ and cleaved DNA (3MQY, dark purple, **Fig. 4E**) reveals a similar rotation and position of the SP (center and right, **Fig. 4C**, **Fig. 4D-E**), with displacements of ∼0.8-0.9 Å (**Table S3**). However, differences are observed in the relative placement of the nucleotides around the cleavage site (**Fig. 4D-E**). The positions of site A cations are similar (0.9 Å and 0.7 Å apart, respectively, **Table S3**) in 3DVO and 3MQY, however the position of the site B ions are 1.1 Å apart when comparing SgrAI_K242A_/40-1/Mg^2+^ and 3MQY (**Fig. 4E, Table S3**)(Note that 3DVO does not contain a site B metal ion).

### Analysis of the SgrAI_WT_/40-1/Mg^2+^ Structure

To gain insight into the atomic reconfigurations of the enzymatic active site in the stages following catalytic DNA cleavage, we next compared the filamentous SgrAI_WT_/40-1/Mg^2+^ structure to the previously determined structures. According to RMSD analysis (**Table S2**), this structure is most similar to prior filamentous structures. This is not surprising, since the conformations of SgrAI in filamentous and non-filamentous forms differ by large changes as a result of the ∼11° rotation between the two subunits of the SgrAI dimer (**Fig. 1B**)(24). The closest comparison is with the structure of filamentous SgrAI bound to Mg^2+^ and a primary site DNA in which the SP is absent (PDB ID 6OBJ), shown in **Fig. 5A**. However, we observe several shifts: (i) the 3′P is again rotated differently in 6OBJ than the current structure (3′P, **Fig. 5A**), (ii) the 3′OH of the deoxyribose ring of Ade2 (the nucleotide 5’ of the cleavage site) is shifted by 1.2 Å, and (iii) the Mg^2+^ ion in site A is shifted by 1.8 Å between the two structures (**Fig. 5A, Table S3**). We also compared the new structure to that of filamentous SgrAI bound to Ca^2+^ and a primary site DNA containing an intact SP (SgrAI_WT_/40-1/Ca^2+^, PDB ID 7SS5). The major difference between the two structures is the type of metal ion, which either inhibits (Ca^2+^ in 7SS5) or promotes (Mg^2+^, the current structure) DNA cleavage. The comparison between PDB 7SS5 and SgrAI_WT_/40-1/Mg^2+^ is shown in **Fig. 5B**. Notably, we find that the distances between the divalent cations in sites A are 0.9 Å apart, and those in site B are 1.7 Å. Concomitant with the shift in the position of the metal ions, we also observe a large displacement of the SP, as expected given that the structures represent two distinct stages of the enzymatic catalytic cycle (**Fig. 5B, Table S3**).

**Figure 5.**
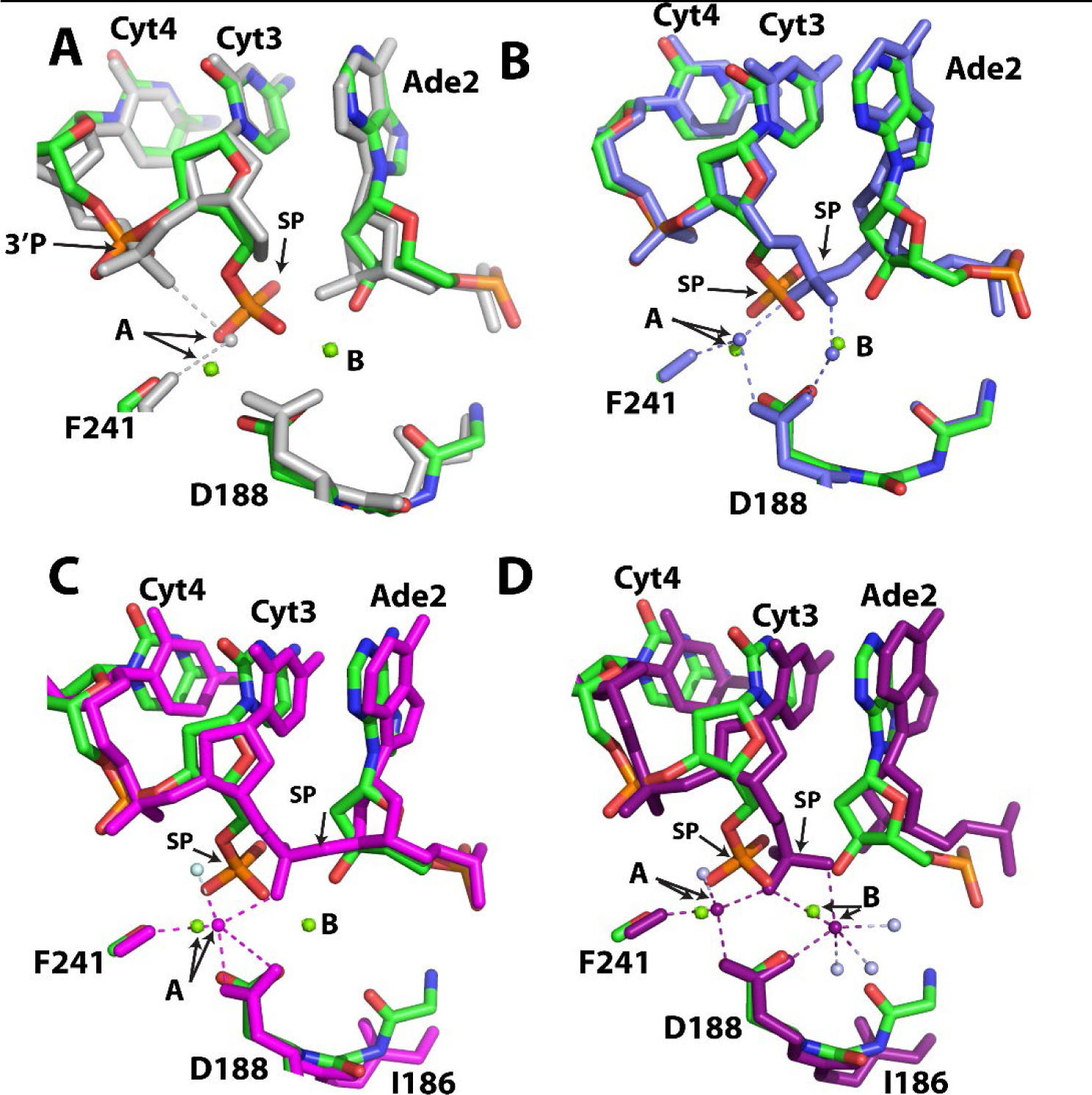
Comparison of SgrAI_WT_/40-1/Mg^2+^ with other SgrAI structures. **A.** All atoms of one chain were used in alignments. SgrAI_WT_/40-1/Mg^2+^ is shown in green, red, blue, orange and SgrAI_WT_/PC/Mg^2+^ (PDB ID 6OBJ) is shown in white. PC is a pre-cleaved version of 40-1 missing the phosphate at the cleavage site (i.e. the scissile phosphate or SP). **B.** Superposition of SgrAI_WT_/40-1/Mg^2+^ (green, red, blue, orange) with SgrAI_WT_/40-1/Ca^2+^ (PDB ID 7SS5, slate). **C.** Superposition of SgrAI_WT_/40-1/Mg^2+^ (green, red, blue, orange) with SgrAI_WT_/18-1/Ca^2+^ (PDB ID 3DVO, magenta, water molecule in light blue). **D.** Superposition of SgrAI_WT_/40-1/Mg^2+^ (green, red, blue, orange) with SgrAI_WT_/18-1/Mg^2+^ (PDB ID 3MQY, magenta, water molecules in light blue).

We next compared the structure of SgrAI_WT_/40-1/Mg^2+^ to non-filamentous SgrAI assemblies, which are expected to reveal more significant changes. **Figure 5C** shows a superposition of SgrAI_WT_/40-1/Mg^2+^ with SgrAI_WT_/18-1/Ca^2+^ (PDB ID 3DVO), the latter being the non-filamentous form bound to a primary site DNA and Ca^2+^ ions (25). Because this form of the enzyme is bound with Ca^2+^, the DNA is uncleaved. Only site A is occupied in 3DVO, and the distance between the Ca^2+^ ion in site A of 3DVO and the Mg^2+^ ion in site A of SgrAI_WT_/40-1/Mg^2+^ is 0.8 Å (**Fig. 5C, Table S3**). The superposition shows that the positions of the nucleotides differ more than in comparisons with any of the filamentous forms. The difference in the SP position within the two structures can be attributed to the state of DNA cleavage.

Finally, the structure of non-filamentous SgrAI containing a cleaved primary site DNA and Mg^2+^ (PDB ID 3MQY) (26) represents the same product structure as SgrAI_WT_/40-1/Mg^2+^ but in the non-filamentous state (**Fig. 5D)**. In this comparison, we observe significant shifts in the structure of the DNA, including both the backbone and the bases (**Fig. 5D**). The phosphorus atoms of the SP groups in the two structures are 1.7 Å apart, and the O3’ atoms are 2.2 Å apart. The differences in the position of the divalent cations in sites A and B are 0.7 Å and 1.2 Å, respectively. Coordination to the Mg^2+^ ions also differs in important ways in these two product structures. In the SgrAI_WT_/40-1/Mg^2+^ structure, each Mg^2+^ coordinates to a single unique non-bridging oxygen of the cleaved phosphate (**Fig. 5D**). In 3MQY, one non-bridging oxygen of the SP coordinates to both Mg^2+^ ions in sites A and B, while a second oxygen coordinates site B. Hence the site B Mg^2+^ coordinates to two oxygen atoms of the cleaved SP in 3MQY, but only one in SgrAI_WT_/40-1/Mg^2+^. The position of the SP is shifted toward the expected position of the nucleophile in SgrAI_WT_/40-1/Mg^2+^ (predicted to be coordinated to the site A Mg^2+^), but the SP is closer to the site B Mg^2+^ in 3MQY. In the SgrAI_WT_/40-1/Mg^2+^ structure, the O3’ leaving group of the cleavage reaction is coordinated to the Mg^2+^ ion at site B, but this coordination is absent in 3MQY.

In summary, comparison of the two new structures to those determined previously shows variability in (*i*) the conformation and positioning of the phosphodiester 3′ of the cleavage site (i.e. the 3′P), (*ii*) the conformation of the SP, (*iii*) variability in the positioning of the site A and B metal cations, and (*iv*) variability in the positioning of the nucleotides around the active site. These differences are relevant to understanding the enzymatic cleavage mechanism, as will be discussed below.

### Comparison of SgrAI_K242A_/40-1/Mg^2+^ and SgrAI_WT_/40-1/Mg^2+^

Finally, we compare the two new structures, SgrAI_K242A_/40-1/Mg*^2+^* and SgrAI_WT_/40-1/Mg*^2+^*, directly (**Fig. 6A-B**). The imaged assemblies and the corresponding experimentally derived structures differ only by the presence or absence of the side chain of lysine 242 in SgrAI, which is one of the active site triad residues. SgrAI belongs to the class of endonucleases with the motif composed of two acidic residues, as well as a third residue that is most often a lysine: PD…(D/E)x**K** (43–46). The acidic residues function to coordinate the divalent cations, which are critical for the DNA cleavage reaction, while the role of the lysine has been debated and will be discussed further below. The structures align well, with the largest changes occurring at the SP, which is cleaved in the WT structure, but uncleaved in the K242A mutant structure. The phosphorus atoms are 1.0 Å apart in the superposition (**Table S3**), residing closer to the expected location of the nucleophile in the structure with cleaved DNA. This also allows for the necessary space to occur between the cleaved phosphate and the free O3′ produced following DNA cleavage (**Fig. 6A-B**). No large shifts are observed in the positioning of the nucleotide bases (**Fig. 6A-B**). The position of the site A cation is similar in the two structures (0.7 Å RMSD, **Fig. 6A-B, Table S3**), but the site B cation is considerably shifted by 1.5 Å between the two. This may be caused by the shift of the cleaved phosphate away from the site B binding site and towards the site A binding site, which is expected following nucleophilic attack (**Fig. 6A-B**). In summary, with the exception of the SP, the positioning of the important active site moieties in these two new structures are closest amongst all structural comparisons, and therefore they should better represent the structural states of the filamentous assembly “before” and “after” DNA cleavage.

**Figure 6.**
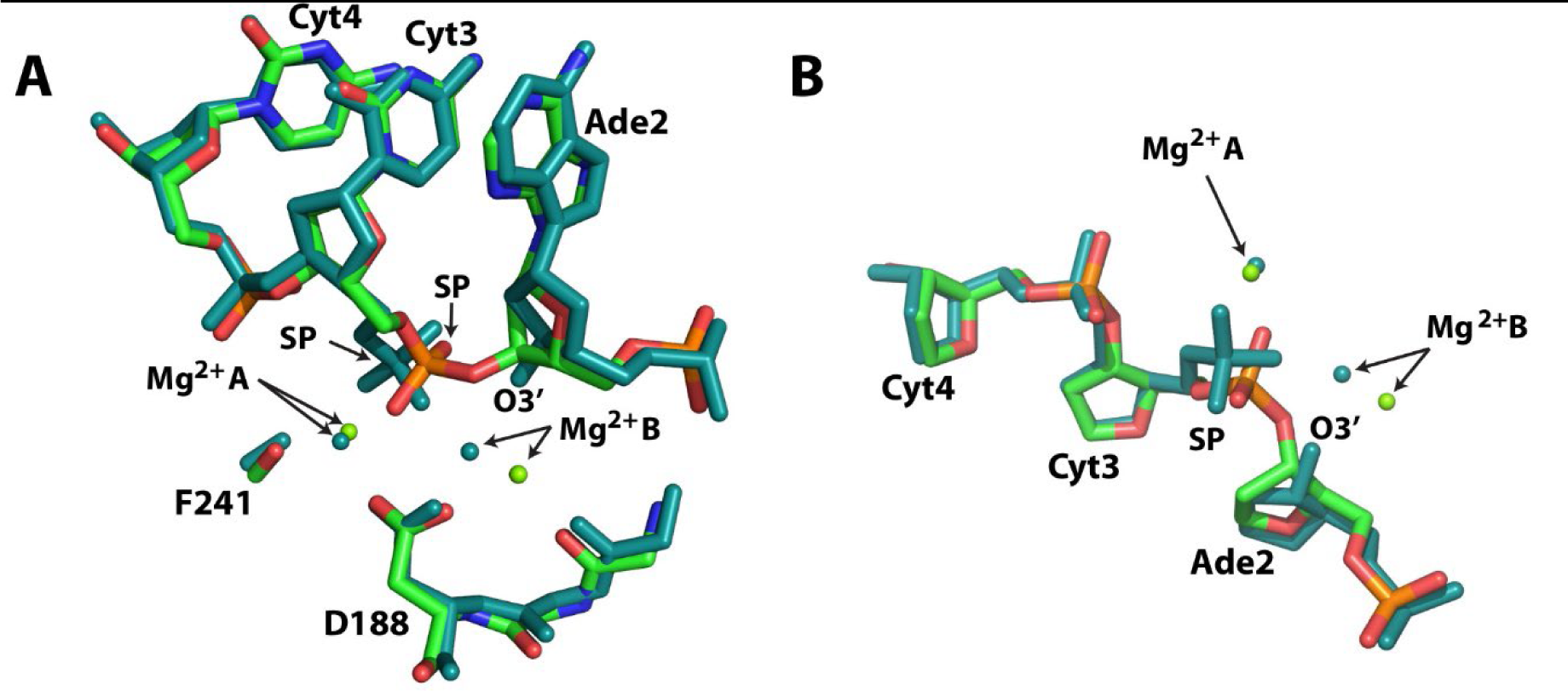
Comparison of SgrAI_K242A_/40-1/Mg^2+^ and SgrAI_WT_/40-1/Mg^2+^ structures. **A.** View of the active site after superposition using all chain A atoms. Wild type SgrAI shown in teal, K242A mutant shown in green, red, blue, and orange. SP, scissile phosphate. **B.** As in A, view from above showing the DNA backbone at the SP and positions of Mg^2+^ ions.

## Discussion

To investigate the binding mode of the biologically relevant metal ion cofactor, as well as the mechanism of filamentation-induced activation of SgrAI, we determined two new structures of filamentous SgrAI bound to primary site DNA and Mg^2+^. Prior structures of filamentous SgrAI were determined using strategies that prevent the DNA cleavage reaction and capture the active site prior to nucleophilic attack, such as metal ion substitution or DNA modification (24,34). While these prior structures provided important insights into the conformational changes induced by filamentation, important questions remained unanswered, which concerned the mechanism of DNA cleavage and the structural alterations that arise from substitutions and modifications used to stall enzymatic cleavage. For example, one structure contained Ca^2+^ in place of Mg^2+^ (PDB ID 7SS5) (34). Mg^2+^ and Ca^2+^ share similar coordination chemistries, and thus Ca^2+^ will often bind to similar sites in proteins but prevent, rather than assist, in the DNA cleavage reaction (35,36). The origin of this effect – inhibition vs. activation – has been the subject of debate and could be related to the larger size of Ca^2+^, to differences in its coordination chemistry, or to its lower ability to polarize coordinated water molecules (33,41,42). Similarly, the structure containing the DNA modification (PDB ID 6OBJ) possessed no phosphodiester bond at the site of DNA cleavage, and hence that structure could not define the binding configuration of this important group (24). The absence of the SP also likely disrupted other important features of the active site, such as occupation of site B by Mg^2+^ (24). One of the two new structures presented herein contains no substitutions or modifications and reveals the active site configuration of the WT enzyme with Mg^2+^ bound, post-DNA cleavage. The other new structure also incorporates Mg^2+^ but employs the K242A point mutation in the active site of SgrAI to stall DNA cleavage. Both structures revealed the anticipated binding of Mg^2+^ in both sites A and B and the expected configuration of the DNA, thereby adding new information to our understanding of the mechanism of DNA cleavage by activated filamentous SgrAI.

### The-Two-Metal-Ion Mechanism

SgrAI is a divalent cation-dependent DNA endonuclease, and analogous to many DNA nucleases, likely uses the two divalent cations in a two-metal-ion cleavage mechanism (**Fig. 1D**). This mechanism was first proposed for alkaline phosphatase (31) and the 3′->5′ exonuclease activity of DNA Pol I from *E. coli* (27), but has since been proposed, with various modifications, to describe the reaction mechanisms of many divalent cation nucleases and other phosphoryl transfer enzymes (28–30,33,47–50). In this mechanism, the two divalent metal cations are coordinated by enzyme moieties such as aspartic or glutamic acid residues and positioned at either side of the scissile phosphodiester bond to be cleaved (**Fig. 1D**). The reaction is thought to occur via an S_N_2(P) type associative/addition-elimination reaction with a true pentacovalent phosphorane intermediate, or alternatively via a concerted reaction mechanism where bond formation and breakage occur simultaneously and the pentacovalent species is instead a transition state (50,51). The cleavage reaction is catalyzed by the enzyme via (*i*) positioning of reactive groups, (*ii*) activation of the nucleophile, (*iii*) stabilization of the transition state, and (*iv*) stabilization of the leaving group (50). These effects are accomplished in part by the divalent cation in site A, which coordinates a water molecule positioned appropriately for nucleophilic attack on the phosphorus atom of the SP. The water is within van der Waals distance (∼3.3 Å) from the phosphate atom, and “in-line” with the leaving group, meaning that the angle between the nucleophile, phosphate atom, and the leaving group (the O3′) is ∼180° (52–54). In addition to positioning, coordination to the site A metal ion also lowers the pKa of the water, such that it is more likely to lose a proton and form the more nucleophilic hydroxide ion (41,42). Both metal ions A and B are positioned to electrostatically stabilize the transition state, which will incur an additional negative charge upon nucleophilic attack. Metal ion B performs the important role of leaving group stabilization via direct coordination or protonation by a metal ion ligated water molecule (or both) (50). Stabilizing the negative charge, which forms on the leaving group in the transition state and after bond cleavage, is thought to be very important to catalytic rate enhancement of this reaction, as this oxygen is not particularly acidic and bearing a negative charge would be highly disfavored (50).

### The role of K242, the active site lysine

Many restriction endonucleases, as well as nucleases containing the restriction endonuclease fold, contain a lysine residue in their active site motif PD…(D/E)x**K** (44). Mutation of this lysine results in greatly diminished rates of DNA cleavage (55–58). Consistent with these observations, our structure of SgrAI containing the K242A mutation showed no evidence of DNA cleavage, despite the presence of the biologically relevant and catalytically competent Mg^2+^ cofactor. While it is clear that the K242 side chain is important to the DNA cleavage activity of SgrAI and similar enzymes, the role it plays in cleavage catalysis remains poorly understood. Since its identification, the active site lysine has been speculated to perform roles such as acting like a general base, positioning of reactive groups, activation of the nucleophile, and stabilization of the transition state (44,57,59–61). Unfortunately, little experimental evidence exists to distinguish among these possible roles. However, a role as a general base seems less likely, because it would require the pKa of the side chain to be significantly reduced below its usual ∼10-11; our structures clearly indicate that the lysine maintains important electrostatic contacts with the negatively charged SP via salt bridge interactions, (**Fig. 7**), an interaction expected to raise its pKa.

**Figure 7.**
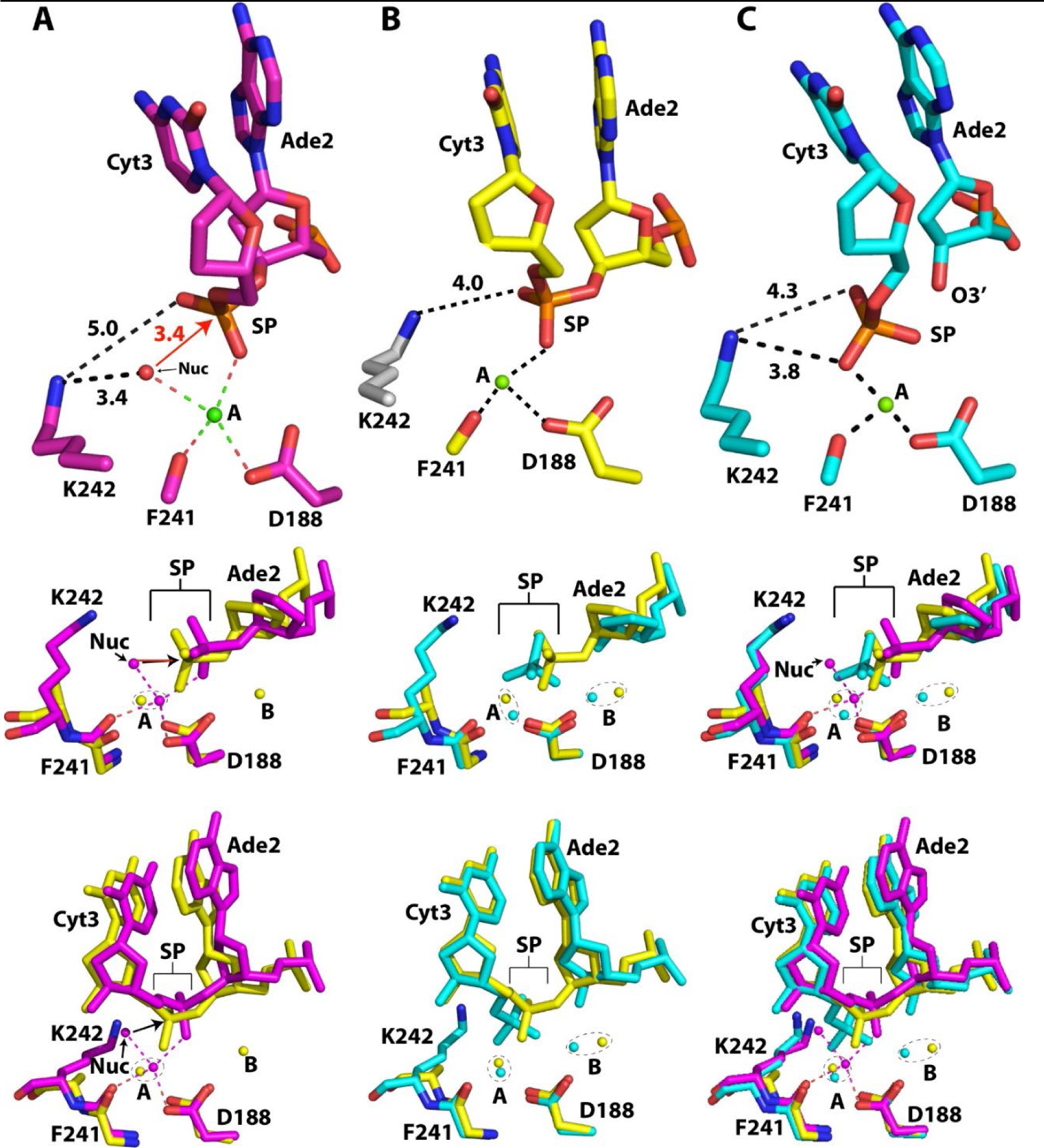
Position of K242 side chain in pre- and post-cleavage structures. **A.** Top panel, active site arrangements in 3DVO. Middle and lower panels, superposition using all atoms of one chain of 3DVO (pink) and SgrAI_K242A_/40-1/Mg^2+^ (yellow). **B.** Top panel, active site arrangement in SgrAI_K242A_/40-1/Mg^2+^. Side chain of K242 (white) is from 3DVO to show its possible position. Middle and lower panels, superposition using all atoms of a single chain of SgrAI_K242A_/40-1/Mg^2+^ (yellow) and SgrAI_WT_/40-1/Mg^2+^ (cyan). **C**. Top panel, active site arrangement in SgrAI_WT_/40-1/Mg^2+^, middle and lower panels, superposition using all atoms of a single chain of 3DVO (pink), SgrAI_K242A_/40-1/Mg^2+^ (yellow) and SgrAI_WT_/40-1/Mg^2+^ (cyan).

**Figure 7** shows the location of the lysine side chain relative to the SP and to the site A metal cation in the active sites of three representative structures. The pre-cleavage state of SgrAI is represented by the structure of non-filamentous SgrAI bound to DNA and Ca^2+^, (PDB ID 3DVO, **Fig. 7A**) as well as the new structure of filamentous SgrAI_K242A_/40-1/Mg^2+^ (**Fig. 7B**). The post-cleavage state of SgrAI is represented by the structure of filamentous SgrAI_WT_/40-1/Mg^2+^ (**Fig. 7C**). Prior to DNA cleavage, the putative water/hydroxide nucleophile (Nuc, upper panel **Fig. 7A**) is coordinated to the site A cation and is also within a salt bridging distance to the terminal amine of K242A (3.4 Å). In the post-cleavage state (upper panel, **Fig. 7C**), the position occupied by the putative nucleophile in **Figure 7A** is now occupied by a nonbridging oxygen of the cleaved SP, implying that this oxygen atom is derived from the nucleophile. The terminal amine of the K242A side chain remains within salt bridging distance with both the nucleophile and the SP, suggesting possible roles of the lysine side chain in catalysis, including positioning the nucleophile and SP for in-line attack, activation of the nucleophile, and stabilization of the transition state. Interestingly, the absence of the K242 side chain results in a shift in the position of the SP towards the space left by the missing side chain (upper panel, **Fig. 7B** and in yellow in the middle panel, **Fig. 7A**), consistent with a role of the side chain in positioning the SP. Water molecules are poorly visible at the resolution of the SgrAI_K242A_/40-1/Mg^2+^ structure, and thus the position of the nucleophile cannot be unambiguosly defined. However, nucleophilic activation could be performed by electrostatic stabilization of the deprotonated state by the positively charged amino group, in other words, via lowering of the pKa of the nucleophilic water by the protonated lysine side chain (61). Although this water is also coordinated to the Mg^2+^ at site A, a Mg^2+^ ligated water has a pKa of 11.4 (42), hence additionally lowering to the pH optimum of the reaction (∼8) would be catalytic (61). In addition to positioning and nucleophile activation, the lysine side chain may also contribute to catalytic rate enhancement via transition state stabilization. The transition state is expected to contain an additional negative charge following nucleophilic attack, which is why the formation of the salt bridge mediated by lysine 242 that is observed in multiple structures (upper panels, **Fig. 7**) would contribute to lowering its energy and thereby increasing the rate of reaction.

### SgrAI and the Two-Metal-Ion Mechanism

SgrAI is one of a growing number of enzymes known to form reversible polymeric filaments. In the case of SgrAI, filamentation results in accelerated DNA cleavage and expanded DNA sequence specificity. We first discovered filamentation by SgrAI in 2010 (19). Since that time, we have determined structures of SgrAI in both filamentous and non-filamentous states to understand the structural origins of the observed filamentation-induced activation and modulation of specificity (23–26,34). The two new structures presented here clarify key outstanding questions regarding the activated DNA cleavage mechanism mediated by SgrAI. For example, we now see for the first time Mg^2+^ occupy site B in a filamentous structure, and we also see the orientations of the scissile phosphate (SP), which are consistent with the two-metal-ion mechanism. In comparison, the prior structure of filamentous SgrAI with site B occupied by Ca^2+^ showed an unexpected rotation of the SP (slate, **Fig. 4B-C**) (34), which may contribute to the failure of Ca^2+^ to stimulate DNA cleavage by SgrAI. **Figure 7** shows the overlay of the two new filamentous SgrAI structures, aligned using all atoms of a single chain of SgrAI. Because fewer differences are seen in these before and after snapshots of the active site, with the exception of the bonds being made and broken, these structures provide a clearer picture of the reaction mechanism.

**Figure 8** summarizes our current working model of the activated DNA cleavage mechanism mediated by SgrAI. First, a non-bridging oxygen of the scissile phosphate, along with a carboxylate oxygen of D188 (and a carbonyl oxygen of F241, not shown) coordinates the Mg^2+^ ion in site A (leftmost panel, **Fig. 8**). In the filamentous conformation, the segment containing I186 shifts and its carbonyl oxygen hydrogen bonds to a water molecule, which stabilizes the binding of a Mg^2+^ ion in site B (second panel, **Fig. 8**). Other ligands of the coordination sphere of the site B Mg^2+^ ion include a carboxylate oxygen of D188 and a non-esterified oxygen of the SP (second panel, **Fig. 8**). The site A Mg^2+^ coordinates a water molecule that will have had its pKa shifted, allowing for greater deprotonation at physiological pH (41,42). The resulting hydroxide is more nucleophilic and is positioned in-line for attack on the phosphorus atom (second panel, **Fig. 8**). Following nucleophilic attack, the phosphorus atom has 5 bonded oxygen atoms, and takes on a trigonal bipyramidal geometry (third panel, **Fig. 8**) (52–54). This species may be a true intermediate, or alternatively a transition state (50). In our model, the site B Mg^2+^ moves closer to the SP at this point, and directly coordinates the O3′ atom, providing a catalytic function in stabilizing the negative charge that forms in the transition state and that will be fully realized upon breakage of the P-O3′ bond. The water molecule, which links the movement of residues in the vicinity of I186 to the creation of site B is positioned to donate a proton to the O3′, which also stabilizes this leaving group.

**Figure 8.**
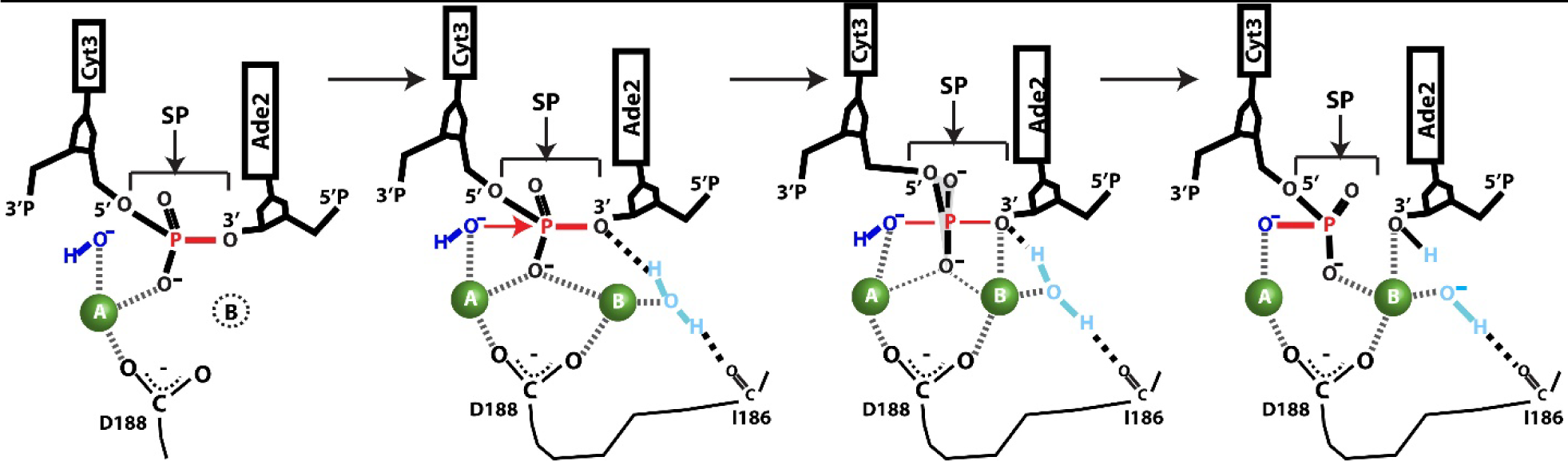
Updated model of active DNA cleavage by SgrAI.

We note two important differences in the current SgrAI structures and the general two-metal-ion mechanism. Although we propose direct coordination by metal ion B to the O3′ and nonbridging oxygen of the SP prior to nucleophilic attack (second and third panels, **Fig. 8**), these interactions are not observed in the SgrAI_K242A_/40-1/Mg^2+^ structure (the distance between the site B ion and the nonbridging oxygen is 4.2 Å, and between the site B ion and the O3’ is 3.1 Å). In the SgrAI_K242A_/40-1/Mg^2+^ structure, the SP has moved towards the site A metal ion (relative to 3DVO, middle panel, **Fig. 7A**), possibly due to the K242A mutation, hence the SP is too distant to coordinate the site B ion. However, direct coordination of the O3′ may not be necessary, and protonation from a site B water may be sufficient to lower the barrier to bond breakage (50,62). Alternatively, coordination of the O3′ may occur as the reaction proceeds (see **Fig. 8**, transition from second to third panel), consistent with its observed coordination to site B in the SgrAI_WT_/40-1/Mg^2+^ structure. In addition, the distances between the carboxylate atoms of D188 and the Mg^2+^ ions in both sites A and B in the SgrAI_K242A_/40-1/Mg^2+^ structure are relatively long for direct coordination (2.9 and 2.8 Å, respectively, **Fig. 3B**). This may be a consequence of coordinate error in structure refinement, or to the observed shift in the position of the SP. Finally, we note that while the site A cation appears to shift very little in our structural comparisons, the site B cation takes on positions up to nearly 2 Å apart suggesting greater flexibility and ability to shift during the reaction pathway.

### Conclusions

In recent years, it has become clear that many important enzymes form filaments with altered properties such as increased or decreased enzymatic rates, cooperativity, or sensitivity to allosteric effectors (1,63–65). Altering substrate specificity is less common (66), but exemplified by SgrAI, which forms filaments with accelerated DNA cleavage activity and expanded DNA sequence specificity (6). The studies described herein add to our growing understanding of the mechanisms by which filamentation activates the DNA cleavage properties of SgrAI. The new structures contain the biologically relevant cofactor Mg^2+^, thus establishing better models in comparison to those derived from prior studies that used metal ion substitutions. The mutant structure (SgrAI_K242A_/40-1/Mg^2+^) contains an active site lysine mutation to stall the DNA cleavage reaction and hence may contain structural changes that are responsible for the absence of catalytic activity. However, this structure also shows occupation of both sites A and B by Mg^2+^ in a pre-cleavage structure. The product structure (SgrAI_WT_/40-1/Mg^2+^) shows a configuration consistent with the structure immediately following DNA cleavage, with interactions among the important residues bridging both metal ions and the active site lysine. Combined with previous structures of filamentous and non-filamentous SgrAI, a clearer picture of the activated DNA cleavage mechanism has emerged. Future studies will be aimed at understanding how filamentation also expands the DNA sequence specificity of SgrAI, which may derive from DNA structure and energetics, as well as disorder-to-order in segments of the SgrAI enzyme (6,34).

## Methods

### Protein preparation

WT SgrAI enzyme was prepared as previously described (20). The expression vector for the K242A mutant of SgrAI was prepared using a commercial source, but because this mutant form had more limited solubility, a second mutation was introduced, L336K. This mutation is far from both the DNA binding site and the interfaces within the filament. Both enzymes were expressed and purified similarly. Briefly, SgrAI enzymes were expressed in BL21 (DE3) *E. coli* (which also contain a constitutive expression system for the methyltransferase enzyme MspI.M) overnight at 18°C. Cells were lysed in lysis buffer (50 mM sodium phosphate buffer, pH 8 at room temperature, 800 mM NaCl, 1 mM 2-mercaptoethanol and 1 mM PMSF) by sonication (using a Fisher Scientific Sonic Dismembrator with a Qsonica Ultrasonic Sonicator converter, Model CL-334) for 3-5 cycles, 5 sec on, 5 sec off, 40% amplitude, and centrifuged for 30 min at 30,000xg to remove cell debris. SgrAI enzymes were purified by chromatography. First, a HisTrap^TM^ FF 5mL (Cytiva^TM^) column was used for an affinity purification step. Protein was bound, washed with lysis buffer and high salt buffer (50 mM potassium phosphate pH 8.0, 2 M NaCl and 1 mM 2-mercaptoethanol) and eluted from the column with a buffer containing 50 mM potassium phosphate pH 8.0, 800 mM NaCl, 1 mM 2-mercaptoethanol and 10-250 mM increasing concentrations of imidazole. Selected fractions, verified through SDS-PAGE, were concentrated using Amicon® Ultra Centrifugal filters (10 kDa cutoff), injected into a Superdex^TM^ 200 Increase GL 10/300 size exclusion column (Cytiva^TM^) and eluted using a buffer containing 25 mM Tris pH 8.0, 150 mM NaCl and 1 mM TCEP. Purified SgrAI enzymes were concentrated and stored in single use aliquots at −80°C in buffer containing 50% glycerol. Enzyme purity was assessed using Coomassie blue staining on SDS-PAGE and assessed to at least 99% purity. Fractions were frozen for subsequent use.

### DNA preparation

Synthetic oligonucleotides purified via C18 reverse phase HPLC were obtained commercially (Sigma-Genosys, Inc.). The concentration was measured spectrophotometrically, with an extinction coefficient calculated from standard values for the nucleotides (67). Equimolar quantities of complementary DNA were annealed by heating to 90°C for 10 minutes at a concentration of 1 mM, followed by slow cooling to room temperature. The sequence of the DNA used in SgrAI/DNA preparations is shown below (red indicates the SgrAI primary recognition sequence):

**40-1-top 5’-GATGCGTGGGTCTTCACACCGGTGGATGCGTGGGTCTTCA-3’**

**40-1-bot 3’-CTACGCACCCAGAAGTGTGGCCACCTACGCACCCAGAAGT-5’**

### Sample preparation for cryo electron microscopy

Both WT SgrAI and K242A mutant SgrAI protein from frozen aliquoted fractions were thawed on ice. The proteins were then subjected to one round of gel filtration chromatography individually with a Superdex 200 Increase 10/300 GL (GE) size-exclusion column, equilibrated in a buffer containing 25 mM Tris-HCl pH 8.0, 150 mM NaCl, 1 mM TCEP, prior to filamentous complex formation. The peak fractions were analyzed by SDS-PAGE gel. The pure fractions from each protein were pooled and concentrated using an Amicon 5 mL 10,000 MWCO centrifugal concentrator (Millipore Sigma, Inc.) for downstream filamentous complex formation.

For SgrAI_WT_/40-1/Mg^2^ filaments, the assembly was made by mixing 30 ul of 3.3 µM SgrAI protein, 1.6 ul of 500 µM 40-1 dsDNA dissolved in water, 3.0 ul of 100 mM MgCl_2_, and the mixture was incubated at room temperature for 1.5 hours. The assembly was centrifuged at 12,000 rpm for 1 minute to remove large aggregates prior to deposition on grids. The Cryo-EM grids were made by applying the centrifuged assembly to R1.2/1.3 gold UltrAufoil grids, Au 300 mesh (Quantifoil), then freezing using a manual plunger in the cold room at 4°C, 90% humidity. For SgrAI_K242A_/40-1/Mg^2+^ filaments, the assembly was made by mixing 30 ul of 12.0 µM SgrAI, 5.8 ul of 500 µM 40-1 dsDNA dissolved in water, 3.1 ul of 100 mM MgCl_2_ and incubated at room temperature for 30 minutes. The assembly was centrifuged at 12,000 rpm for 1 minute to remove large aggregates and then diluted 10-fold using assembly buffer before applying onto graphene grids. The graphene grids were made in-house by depositing a thin layer of graphene over R1.2/1.3 gold UltrAufoil grids, Au 300 mesh (Quantifoil). The Cryo-EM grids were prepared for SgrAI_K242A_/40-1/Mg^2+^ filaments using the same procedures as those used for preparing grids for SgrAI_WT_/40-1/Mg^2+^ filaments. These grids were clipped and then subsequently stored in liquid nitrogen for data acquisition.

### Cryo-electron microscopy data collection

For both SgrAI_K242A_/40-1/Mg^2+^ and SgrAI_WT_/40-1/Mg^2+^ datasets, the movie frames were collected using a Titan Krios transmission electron microscope (Thermo Fisher Scientific) operating at 300 KeV. The data collection was performed in an automated manner using the Leginon software (68,69). For SgrAI_K242A_/40-1/Mg^2+^ dataset, a K3 quantum director (Gatan) with a GIF (Gatan Imaging Filter) BioQuantum energy filter with a slit width of 20 eV was used to record the movies at a magnification of 105,000×, corresponding to a pixel size of 0.83 Å/pixel in nanoprobe EF-TEM mode. The total fluence was 42.8 e^-^/Å^2^ at a rate of 5.1 e^-^/pix/sec. For SgrAI_WT_/40-1/Mg^2+^ dataset, a K2 summit director (Gatan) was used to record the movies composed of 50 frames in counting mode over 4 s (80 ms per frame) at a magnification of 165,000×, corresponding to a pixel size of 0.83 Å/pixel in microprobe EF-TEM mode. The total fluence was 29.6 e^-^/Å^2^ at a rate of 5.1 e^-^/pix/sec. All imaging parameters are summarized in **Table S1**.

### Cryo-EM image analysis

The same workflow was used to process both SgrAI_K242A_/40-1/Mg^2+^ and SgrAI_WT_/40-1/Mg^2+^ datasets. For each dataset, the movie frames were imported to Relion to perform dose-weighted motion-correction on 6 by 6 patch squares and using a B-factor of 100 (70). The motion corrected micrographs were imported into cryoSPARC (71) for downstream data processing. A 2D template was generated after an initial round of particle picking using Blob Picker and 2D classification in cryoSPARC. Subsequently, the 2D template was used for template-based particle selection in cryoSPARC. The particles were extracted with a box size of 320 pixels after inspection of particle picking. Reference-free 2D classification was used to identify filamentous particles, and after each round of 2D classification, the best 2D class averages were selected based on the appearance of filamentous particles containing good features. When several iterations of 2D classification were performed, the best 2D classes were selected to generate an *ab-initio* reconstruction using three classes as input. The best *ab-initio* with features consistent with filamentous SgrAI was then subjected to homogeneous helical refinement in cryoSPARC with parameters described in **Table S1**. Following homogeneous helical refinement, a heterogeneous refinement was performed using volume inputs from *ab-initio* reconstruction and homogeneous helical refinement. The best volume and its corresponding particles from heterogeneous refinement was selected to perform a homogeneous helical refinement to improve the map. We continue to repeat this procedure, consisting of iterative homogeneous and heterogeneous refinement, until the map resolution and quality does not improve. The best map from the last heterogeneous refinement was subjected a homogeneous helical refinement, followed by one round of per-particle CTF refinement. At this point, the particles were imported into Relion for particle polishing (70,72). Particle polishing was performed in Relion using default parameters. Subsequently, the polished particles were imported back to cryoSPARC to perform per-particle CTF refinement, followed by a homogeneous helical refinement. Iterative Bayesian polishing in Relion and homogeneous refinement in cryoSPARC was repeated until no further improvements in map resolution were observed, as assessed using the Fourier shell correlation. To segment an SgrAI dimer from the helical map, a mask was generated using EMAN2 (73,74) and Chimera (75). The local resolution was calculated in cryoSPARC. The 3D FSC (76) was obtained using the 3D FSC server (3dfsc.salk.edu) and the sampling compensation function (SCF) (77) was calculated using the graphical user interface tool (78). Image analysis results are shown in **Figures S1-S2** and summarized in **Table S1**.

### Atomic model refinement filamentous SgrAI structures from cryo-EM maps

We used an atomic model derived from the previously determined 2.7 Å cryo-EM map of filamentous SgrAI containing Ca^2+^ and an intact SP (PDB 7SS5(34)), to build and refine the model of SgrAI_K242A_/40-1/Mg^2+^ and SgrAI_WT_/40-1/Mg^2+^. The models were built/adjusted (including the SP, the metal ions, and the addition of water molecules) in Coot(79). Subsequently, we performed one round of real-space refinement within the Phenix (80) suite. The models were iteratively adjusted in Coot and refined in Phenix (81), and the statistics were examined using Molprobity (82) until no further improvements were observed. The final models were also evaluated using FSC analysis against the map and using EMRinger (83) to compare the fit of the model backbone into the cryo-EM map. The model statistics showed good geometry and matched the cryo-EM reconstruction (**Figures S1-S2 and Table S1**). Refined coordinates (9BGJ, 9BGI) and maps (EMD-44514, EMD-44513) have been deposited for the SgrAI_K242A_/40-1/Mg^2+^ and SgrAI_WT_/40-1/Mg^2+^, respectively, in the appropriate databases.

### Structural analysis

Structures were aligned using all atoms of a single SgrAI chain using Pymol(84). RMSD were calculated using Pymol(84) and Chimera(75) using all atoms or only alpha carbon atoms, as indicated. Figures were prepared with Pymol(84) and Chimera(75).

## Funding and Additional Information

Research reported in this publication was supported by the National Science Foundation under Grant No. MCB-1934291 (N.C.H. and D.L.), and the University of Arizona Research, Innovation & Impact (RII) and Technology Research Initiative Fund/Improving Health and Access and Workforce Development (N.C.H.). D.L. also acknowledges support by the National Science Foundation (MCB 2048095) and the support of the Hearst Foundations developmental chair. A.R.G. acknowledges the support of the Pathways in Biological Sciences (PiBS) NIH T32 Training grant (T32 GM133351) and the Mary K. Chapman Foundation.

---

**Figure S1.**
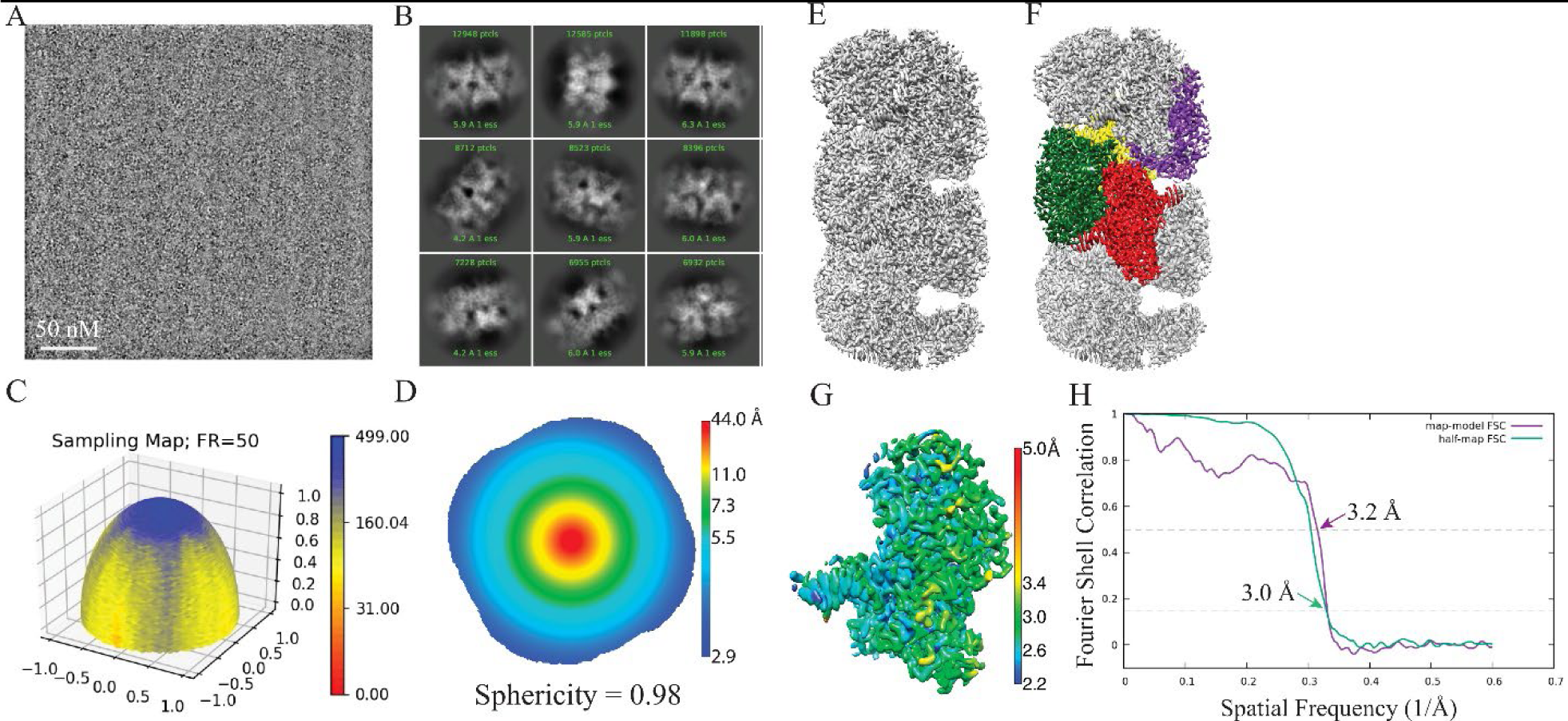
Cryo-EM data and validation of SgrAI_K242A_/40-1/Mg^2+^. (**A**) A micrograph showing the particle distribution on an ultrAufoil grid overlaid with graphene. (**B**) Example 2D averages from the data used in the final reconstruction map. (**C**) Surface sampling plot derived from the Euler angle distribution, calculated with a Fourier radius set to 50 voxels. The sampling compensation factor (SCF) is indicated in **Table S1** (77,78). (**D**) Central slice through the 3DFSC (76,85) colored by resolution. (**E**) The full cryo-EM map. (**F**) The full cryo-EM map with 4 DBDs colored as in **Fig. 1A**. (**G**) A segmented DBD from the map, colored by local resolution. (**H**) Fourier shell correlation (FSC) curves derived from half-map and map-to-model reconstructions, with FSC cutoffs 0.143 and 0.5 indicated, respectively, as well as nominal resolution values.

**Figure S2.**
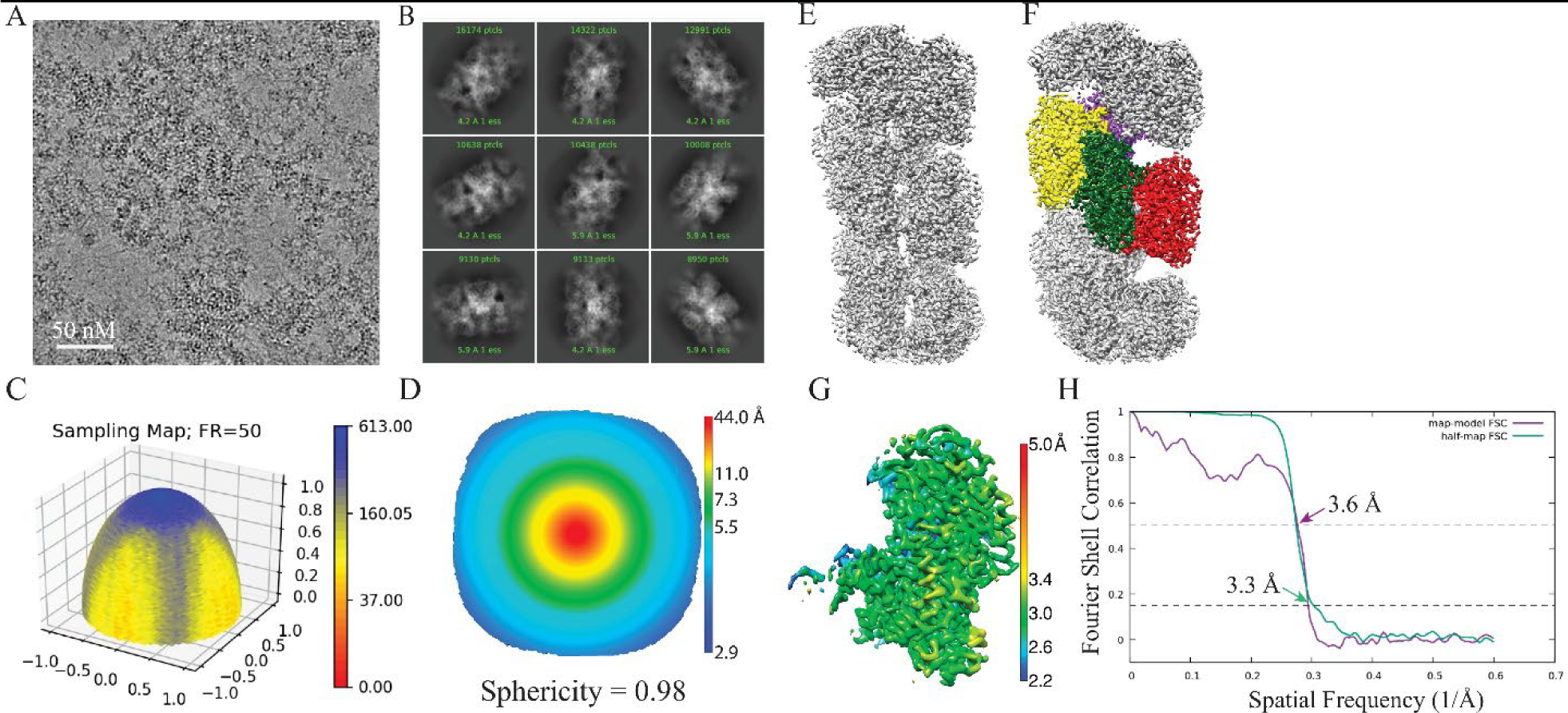
Cryo-EM reconstruction validation of SgrAI_WT_/40-1/Mg^2+^. (**A**) A micrograph showing the particle distribution on an ultrAufoil grid overlaid with graphene. (**B**) Example 2D averages from the data used in the final reconstruction map. (**C**) Surface sampling plot derived from the Euler angle distribution, calculated with a Fourier radius set to 50 voxels. The sampling compensation factor (SCF) is indicated in **Table S1** (77,78). (**D**) Central slice through the 3DFSC (76,85) colored by resolution. (**E**) The full cryo-EM map. (**F**) The full cryo-EM map with 4 DBDs colored as in **Fig. 1A**. (**G**) A segmented DBD from the map, colored by local resolution. (**H**) Fourier shell correlation (FSC) curves derived from half-map and map-to-model reconstructions, with FSC cutoffs 0.143 and 0.5 indicated, respectively, as well as nominal resolution values.

**Table S1.** Cryo-EM Data and Model Refinement Statistics.

| **EM Data Collection/Processing** |  |  |
| --- | --- | --- |
|  | wtSgrAI/40-1/Mg<sup>2+</sup> | K242A/40-1/Mg<sup>2+</sup> |
| Submission codes | 9BGI, EM-44513 | 9BGJ, EMD-44514 |
| Microscope | FEI Titan Krios | FEI Titan Krios |
| Voltage (kV) | 300 | 300 |
| Camera | Gatan K2 Summit | K3 quantum |
| Magnification | 165,000 | 105,000 |
| Nominal defocus range (μm) | 0.8-2.0 | 1.0-2.5 |
| Exposure time (s) | 4 | 750 |
| No. of frames | 50 | 50 |
| Frame rate (frames per second) | 12.5 | 66.7 |
| Total fluence (e-/Å<sup>2</sup>) | 29.6 | 42.8 |
| Fluence rate (e-/pixel/s) | 5.1 | 39.3 |
| Pixel size (Å) | 0.83 | 0.83 |
| No. of movies | 2,704 | 3,488 |
| Total extracted particles | 821,410 | 2,768,698 |
| No. of particles in final map | 202,664 | 357,489 |
| Symmetry | Helical | Helical |
| Helical parameters (rise, Å / twist, °) | 21.3 / -86.2 | 20.9/-86.2 |
| Resolution, Fourier shell correlation 0.143 (Å) | 3.05 | 3.34 |
| Local resolution range (Å) | 1.78 – 6.06 | 1.79 – 5.30 |
| Directional resolution range from 3D FSC (Å) | 2.89 – 3.16 | 3.12 – 3.59 |
| Sampling Compensation Factor (SCF) | 0.780 | 0.738 |
| Map-sharpening B factor (Å<sup>2</sup>) | 87.9 | 133.2 |
| **Atomic model statistics** |  |  |
| Model resolution (Å) | 3.1 | 3.5 |
| FSC ½ threshold |  |  |
| Model composition |  |  |
| Non-hydrogen atoms | 6511 | 6359 |
| Protein residues | 676 | 672 |
| Nucleotides | 48 | 51 |
| Waters | 177 | 9 |
| Ligands (Mg<sup>2+</sup>) | 4 | 2 |
| R.m.s. deviations |  |  |
| Bond lengths (Å) | 0.004 | 0.003 |
| Bond angles (°) | 0.518 | 0.536 |
| Validation |  |  |
| MolProbity score | 1.73 | 2.43 |
| Clashscore | 4.85 | 9.54 |
| Poor rotamers (%) | 1.77 | 4.11 |
| Ramachandran plot |  |  |
| Favored (%) | 95.83 | 92.79 |
| Allowed (%) | 4.17 | 7.21 |
| Disallowed (%) | 0 | 0 |
| CaBLAM outliers (%) | 1.96 | 3.66 |
| Cis proline (%) | 0 | 0 |
| Twisted proline (%) | 0 | 0 |
| Cβ outliers (%) | 0 | 0 |
| Average B-factors (min/max/mean, Å<sup>2</sup>) |  |  |
| Protein | 12.9/145.7/54.3 | 31.4/164.8/78.9 |
| Nucleotide | 8.8/154.3/52.9 | 29.8/209.2/117.4 |
| Ligand | 33.4/44.6/39.3 | 46.4/50.4/48.4 |
| Water | 15.7/87.5/40.0 | 28.9/66.25/54.0 |

**Table S2.** RMSD between structures of SgrAI bound to DNA*.

|  | Using Both Chains of the SgrAI Dimer |  |  |  |  | Using a Single Chain of the SgrAI Dimer |  |  |  |  |
| --- | --- | --- | --- | --- | --- | --- | --- | --- | --- | --- |
|  | SgrAI<sub>WT</sub>/<br>40-1/Mg<sup>2+</sup> | SgrAI<sub>K242A</sub>/<br>40-1/Mg<sup>2+</sup> | 7SS5 | 3DVO | 3MQY | SgrAI<sub>WT</sub>/<br>40-1/Mg<sup>2+</sup> | SgrAI<sub>K242A</sub>/<br>40-1/Mg<sup>2+</sup> | 7SS5 | 3DVO | 3MQY |
| SgrAI<sub>WT</sub>/<br>40-1/Mg<sup>2+</sup> | - | 0.56<br>(0.61) | **0.25**<br>(0.30) | 1.90<br>(1.99) | 1.93<br>(2.07) | - | 0.47<br>(0.54) | **0.22**<br>(0.27) | 0.59<br>(0.70) | 0.68<br>(0.80) |
| SgrAI<sub>K242A</sub>/<br>40-1/Mg<sup>2+</sup> | 0.56 (0.60) | - | 0.56<br>(0.60) | 1.63<br>(1.84) | 1.73<br>(1.92) | 0.47 (0.51) | - | 0.44<br>(0.48) | 0.54<br>(0.66) | 0.62<br>(0.72) |

\*Alpha carbons used in RMSD (all atoms in parentheses)

**Table S3.**
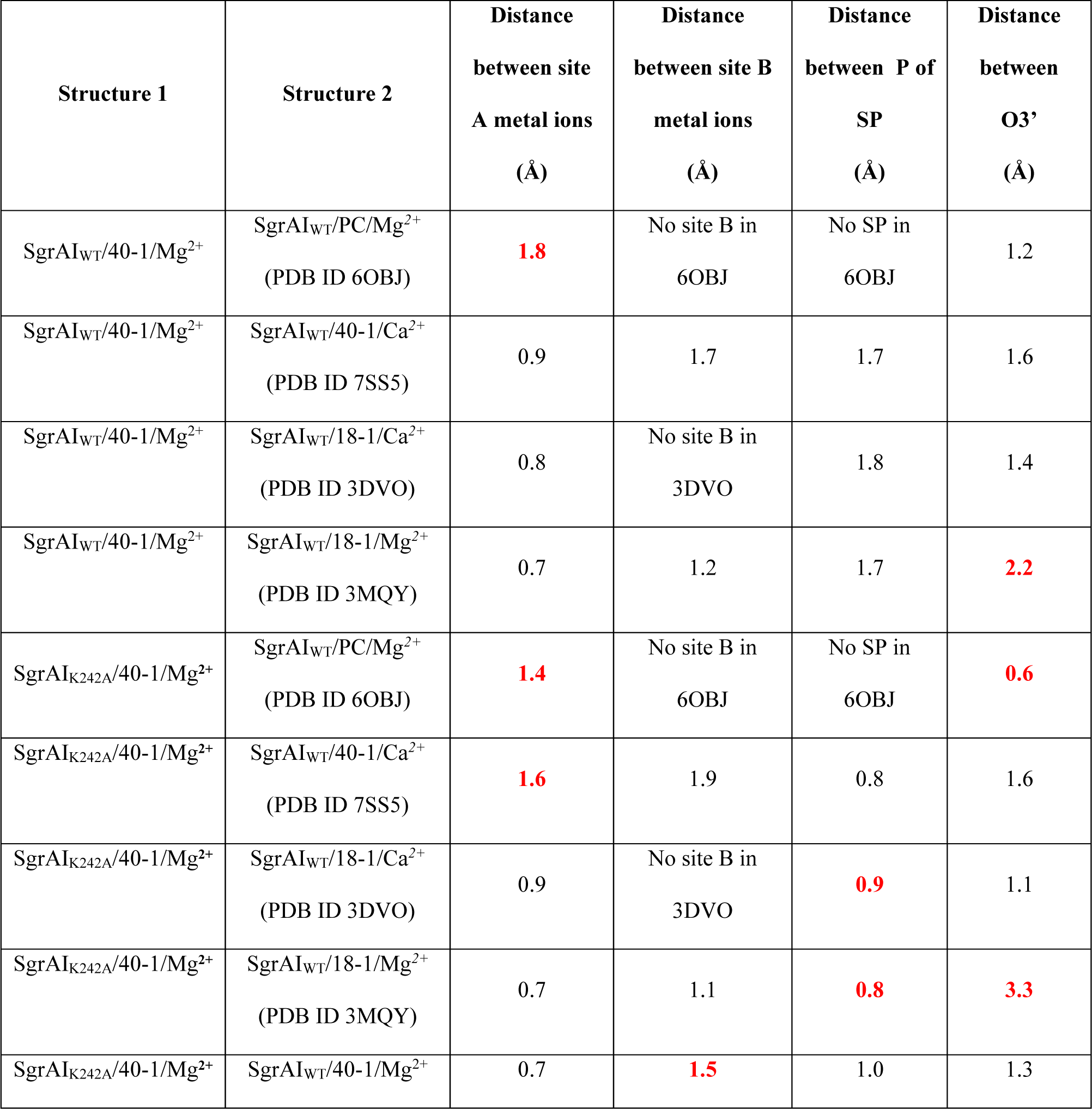
Distances between key atoms in structural superpositions (all atoms of one SgrAI chain used in the superpositions).

